# Identification of HIV-Associated Gene Expression Biomarkers Using Machine Learning and Interpretable Artificial Intelligence

**DOI:** 10.1101/2025.05.08.652807

**Authors:** Sultan Mahmud, Faruk Hossain

## Abstract

Despite advances in antiretroviral therapy (ART), the early and accurate diagnosis of Human Immunodeficiency Virus (HIV) infection remains a significant public health challenge. Traditional biomarkers, such as CD4+ T cell counts and viral load, are limited in capturing the complex biological mechanisms underlying HIV pathogenesis. This study proposes a machine learning (ML) and interpretable artificial intelligence (AI) framework to identify transcriptomic biomarkers for HIV diagnosis using immune cell–specific gene expression data.

We utilized the GSE6740 dataset, which includes microarray profiles from CD4+ and CD8+ T cells of HIV-positive (treatment-naïve) and HIV-negative individuals. Feature selection was performed through differential gene expression analysis, weighted gene co-expression network analysis (WGCNA), and protein–protein interaction (PPI) network construction. Hub genes identified through these methods were used to train six supervised ML classifiers.

The best-performing model was selected based on cross-validated metrics, including accuracy, Kappa, ROC, sensitivity, and specificity, and further interpreted using SHapley Additive exPlanations (SHAP) values. Seven co-expression modules were identified, with the red and green modules showing strong positive and negative correlations with HIV status, respectively. From the intersection of WGCNA modules, differentially expressed genes (DEGs), and PPI networks, ten hub genes were prioritized.

Among the trained models, the regularized regression model (GLMNET) demonstrated the highest diagnostic performance (ROC = 0.97, accuracy = 91%). SHAP analysis highlighted GBP1, ISG15, OAS2, OAS1, and DDX60 as the most influential genes contributing to model predictions, thereby enhancing interpretability and biological relevance.

By integrating transcriptomic profiling with interpretable ML, this study identifies novel gene-based biomarkers for HIV diagnosis and underscores the potential of explainable AI in advancing precision medicine approaches for infectious diseases.

## Introduction

Human Immunodeficiency Virus (HIV) continues to pose a critical global public health challenge, with an estimated 39 million people living with the virus as of 2023 [1]. Despite significant advances in antiretroviral therapy (ART) that have markedly improved the survival and quality of life for people living with HIV (PLWH), substantial challenges remain in the areas of early diagnosis, personalized treatment, and long-term disease monitoring [2]. Current clinical practices primarily rely on biomarkers such as CD4+ T cell counts and plasma viral load to assess disease progression and treatment response. However, these conventional markers often fall short in capturing the complex and dynamic host–virus interactions, and they may not fully reflect the underlying biological processes involved in HIV pathogenesis [3, 4].

In recent years, gene expression profiling has emerged as a promising tool for elucidating host responses to HIV infection. Studies have demonstrated that HIV triggers distinct transcriptional changes associated with immune activation, inflammation, and apoptosis [5-7]. However, many of these investigations have employed traditional statistical approaches that focus on a limited set of genes, which may hinder broader applicability and biological interpretability [8]. Furthermore, the integration of immune-specific transcriptomic data into robust diagnostic frameworks remains limited.

The growing field of machine learning (ML) and interpretable artificial intelligence (AI) presents a promising opportunity to overcome these limitations. ML techniques excel at modeling complex, high-dimensional data and can detect subtle patterns in gene expression that are indicative of disease states [9, 10]. Furthermore, the application of explainable AI methods such as SHapley Additive exPlanations (SHAP) allows researchers to interpret and quantify the impact of individual genes on model predictions, thereby enhancing both the transparency and biological interpretability of diagnostic models [11].

In this study, we propose an integrated ML and interpretable AI framework for identifying diagnostic biomarkers of HIV infection using immune-cell-specific transcriptomic data. We leveraged the publicly available GSE6740 dataset, which includes microarray gene expression profiles from CD4+ and CD8+ T cells of both HIV-positive (treatment-naïve) and HIV-negative individuals [12]. Our analytical pipeline incorporated differential gene expression analysis, weighted gene co-expression network analysis (WGCNA), and protein–protein interaction (PPI) network construction to identify and prioritize HIV-related genes. The most influential hub genes were then used to train a range of supervised ML models—logistic regression, support vector machines (SVM), k-nearest neighbors (kNN), decision trees, random forest (RF), and GLMNET. To ensure interpretability, SHAP values were computed to elucidate the contribution of each feature to model predictions.

This study makes the following novel contributions:

- It integrates WGCNA, differential expression, and PPI network analysis for robust feature selection
- It applies multiple ML algorithms to diagnose HIV infection based on transcriptomic data.
- It incorporates SHAP analysis to ensure transparency and interpretability of predictive models.

By combining computational modeling with biologically informed feature selection, this study aims to advance the development of accurate, interpretable, and clinically applicable diagnostic tools for HIV.

## Method

### Study Design and Data Sources

This study employed a retrospective design to identify HIV-associated gene expression biomarkers by integrating machine learning and interpretable artificial intelligence (AI) techniques. Microarray gene expression data were obtained from the Gene Expression Omnibus (GEO) database. Specifically, the GSE6740 dataset [12], which includes both HIV-positive (case) and HIV-negative (control) samples along with relevant clinical metadata (e.g., age, treatment status, and duration of infection), was used for model development and training.

### Data Preprocessing

The data were transposed to ensure that rows represented samples and columns represented genes, as required by the WGCNA framework. To reduce noise, a pre-filtering step was applied in which genes with zero variance were removed, and the top 11% of genes with the highest variance were retained. As shown in the gene variance curve (S1 Fig 1), the variance sharply drops and flattens beyond this threshold. A log2(TPM + 1) transformation was applied to normalize gene expression levels. Quality control was performed using the goodSamplesGenes function from the WGCNA package to retain only high-quality genes and samples. Phenotypic trait data-including gender, age, infection duration, and HIV status-were extracted and cleaned using regular expressions and string processing functions.

### Differential Gene Expression Analysis

Differentially expressed genes (DEGs) between HIV-positive and HIV-negative individuals were identified using the limma package in R. Genes with an adjusted p-value < 0.05 and |log_2_ fold change| > 0.2 were considered significantly differentially expressed.

### Weighted Gene Co-expression Network Analysis (WGCNA)

To identify gene modules associated with HIV status, WGCNA was performed using the 11% top most variable genes. Modules showing strong correlation with HIV (positive/negative) status were selected for further analysis. The intersection between the significant WGCNA modules and DEGs was used to identify key hub genes for model development.

### Gene Ontology (GO) Enrichment Analysis

To gain insights into the biological significance of the selected genes, Gene Ontology (GO) enrichment analysis was performed using the enrichGO() function from the clusterProfiler R package. The analysis was conducted with the following parameters: the input genes were mapped using their ENTREZ IDs (keyType = “ENTREZID”) against the org.Hs.eg.db database. GO terms across all three ontologies—biological process (BP), molecular function (MF), and cellular component (CC)—were considered (ont = “ALL”). Multiple testing correction was applied using the Benjamini-Hochberg (BH) method (pAdjustMethod = “BH”), with a p-value cutoff of 0.05 and a q-value cutoff of 0.2. Gene IDs were converted into readable gene symbols to facilitate interpretation.

### KEGG Pathway Enrichment Analysis

Kyoto Encyclopedia of Genes and Genomes (KEGG) pathway enrichment analysis was also performed to identify significantly enriched signaling pathways using the enrichKEGG() function. ENTREZ IDs were used for gene mapping against the hsa organism (Homo sapiens). the analysis was conducted with a p-value cutoff of 0.05. The results included measures such as GeneRatio, background ratio (BgRatio), fold enrichment, p-values, adjusted p-values, and q-values.

### PPI Network Analysis and Hub Gene Screening

Protein–protein interaction (PPI) network analysis was performed using the STRING database (version 12.0) to construct a network based on important overlapping co-expressed genes [13, 14]. The top ten hub genes within the PPI network were identified using the Maximum Clique Centrality (MCC) algorithm implemented in the CytoHubba plugin for Cytoscape (version 3.9.3) [15]. The MCC algorithm ranks genes based on their centrality within the network, enabling the identification of key hub genes. The final results, including the top-ranked hub genes, were visualized using Cytoscape.

### Development and assessment HIV diagnostic model

The dataset contained the expression profiles of selected genes, clinical trait status (infected vs. uninfected), age, and other relevant variables, which were used to build machine learning models. A suite of supervised machine learning algorithms was applied to construct diagnostic models based on these features. The following models were trained using the caret package in R:

- Logistic Regression (Logistic) [16]: A generalized linear model using a binomial link function.
- Regularized Regression (GLMNET) [17]: Elastic net regularization via the glmnet method with preprocessing (centering and scaling).
- Support Vector Machine (SVM) with Radial Kernel [18]: Trained using the svmRadial method, with preprocessing.
- K-Nearest Neighbors (kNN) [19]: Classification based on feature proximity, with preprocessing.
- Decision Tree (DecisionTree) [20]: A recursive partitioning tree implemented via the rpart method.
- Random Forest (RF) [21]: A robust ensemble learning method utilizing multiple decision trees.

Model performance was evaluated using 10-fold cross-validation repeated three times to ensure robust estimation and reduce variability. The evaluation metrics used to compare model performance included accuracy, Kappa statistic, area under the ROC curve (AUC), sensitivity, and specificity [20].

Once the best-performing model was identified based on these metrics, a hyperparameter tuning procedure was conducted to optimize the selected GLMNET model using the caret and glmnet packages in R. The alpha parameter determines the type of regularization (Ridge, Lasso, or Elastic Net), while the lambda parameter controls the strength of the regularization. A range of values for both hyperparameters was explored, with alpha values of 0 (Ridge), 0.5 (Elastic Net), and 1 (Lasso), and lambda ranging from 0.0001 to 1 on a logarithmic scale.

A 10-fold cross-validation procedure was applied to ensure that the model’s performance was robust and generalized well to unseen data. The grid search process identified the best combination of alpha and lambda, optimizing the model based on accuracy..

### Model Interpretation

To enhance model interpretability, SHapley Additive exPlanations (SHAP) values were calculated for the top-performing model (GLMNET). SHAP provides global and local feature importance, offering insights into how each gene and clinical variable contributed to the classification outcome.

### Statistical Analysis and Visualization

All analyses were performed using R (version 4.5) and Python (Jupyter-Notebook version 7.2.2). R packages used included caret, randomForest, e1071, glmnet, WGCNA, ggplot2 and SHAP. Summary statistics were presented in tables, while model performance and interpretability outputs were visualized using bar plots, ROC curves, and SHAP summary plots.

## Results

Our study’s overall workflow is depicted in Fig. 1. The dataset GSE6740 included expression profiles for 22283 genes, presenting the typical high-dimensionality challenge common in genomics research. To effectively manage this complexity and identify strong biomarker candidates for HIV infection, we applied several feature selection algorithms. Following feature selection, the development of AI-based diagnostic models was carried out using the most important (hub) genes. These hub genes were selected based on both their statistical significance and their potential biological relevance to disease progression.

**Fig 1.**
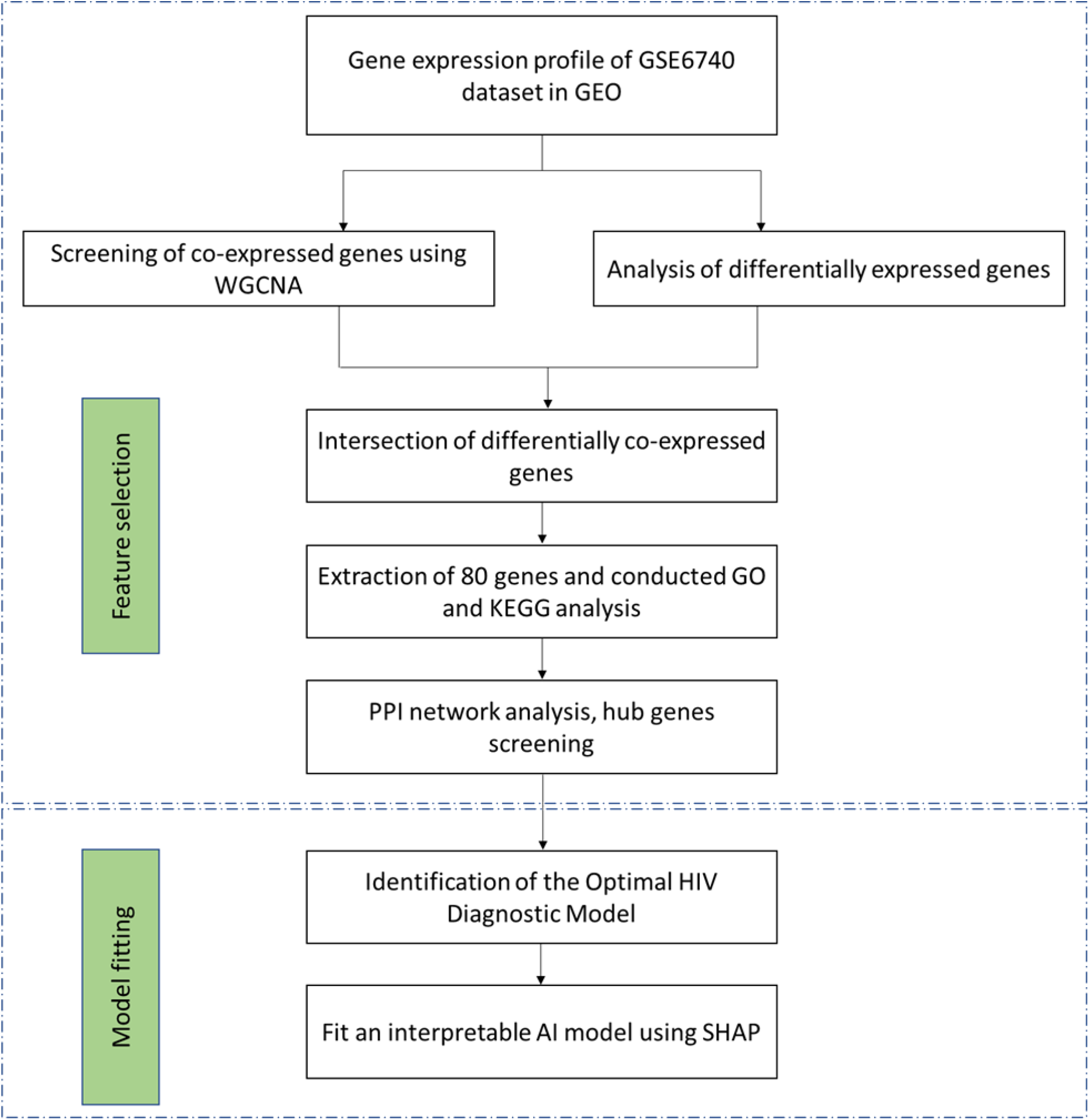
Workflow for identifying and validating diagnostic hub genes for HIV.

### Feature Selection

#### Weighted Gene Co-expression Network Modules

WGCNA analysis was performed on the expression data of 2,451 genes, following the removal of genes with low expression variability (i.e., those outside the top 11% of variance) [22]. A soft-thresholding power of 9 was selected to achieve an optimal balance between scale-free topology fit and network connectivity, based on soft-thresholding power analysis-a critical step in WGCNA to construct a scale-free gene co-expression network (S1 Fig (2(a-b)). According to Fig 2, seven modules (including the gray module) were identified and can be clustered into two group. A heatmap of module–trait relationship was generated to assess the association between each module and the clinical trait, HIV status (HIV+ and HIV−) (Fig 3). This analysis revealed that the red and yellow modules had significant positive association with HIV status (red module: r = 0.54, p = 3e-04; yellow module: r = 0.50, p = 0.001), while the brown and green modules showed significant negative association (brown module: r = -0.4, p = 0.01; green module: r = -0.41, p = 0.009). Focusing on the gene modules, particularly the red strongest disease positive and blue module with the strongest disease negative correlation and most significant p-values, may provide deeper insights into the biological mechanisms of HIV infection and identify potential targets for diagnosis, prognostic assessment, and therapeutic development.

**Fig 2.**
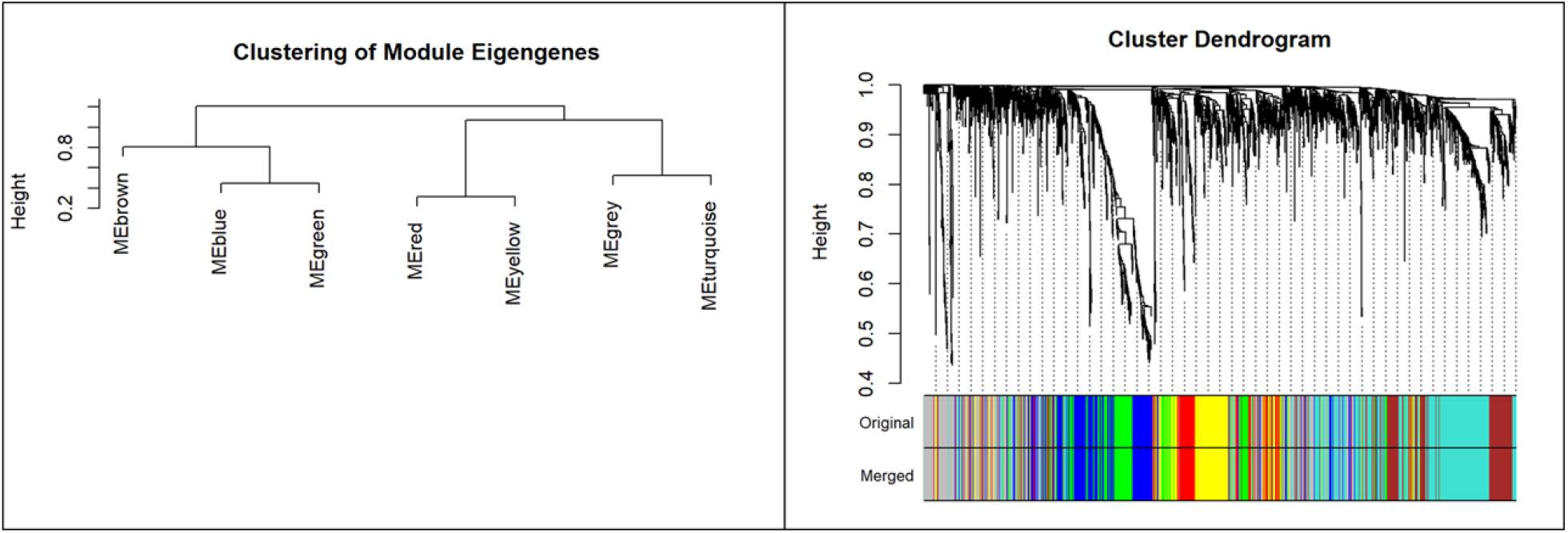
Gene and module clustering dendrogram.

**Fig 3.**
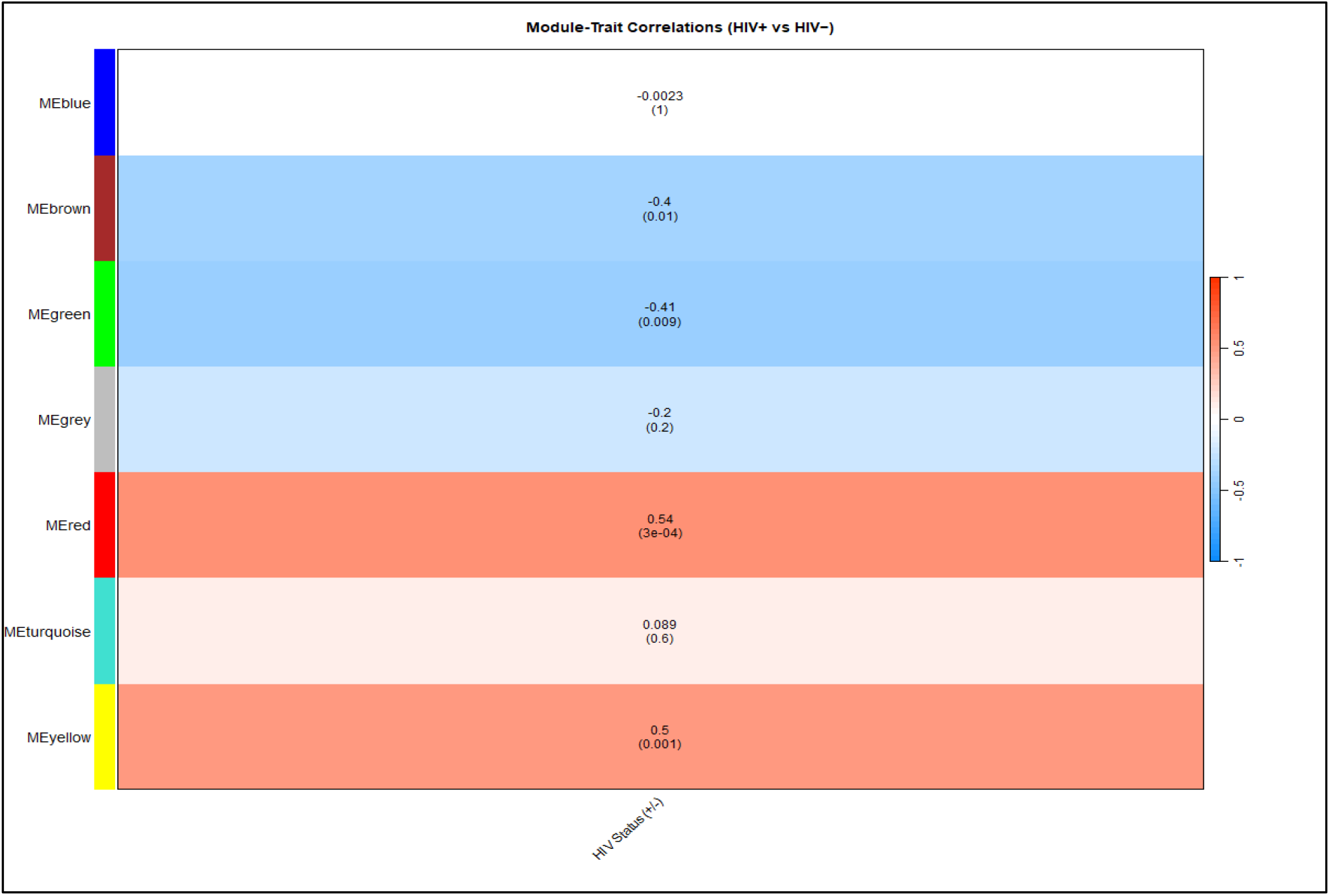
Module trait relationship.

#### Intersection of DEGs and Co-expression Modules

A total of 198 differentially expressed genes (DEGs) were identified in the GSE6740 dataset (|logFC| > 0.2, adjusted p-value < 0.05), including 94 upregulated and 104 downregulated genes (right panel of Fig 4). As illustrated in the Fig, 170 co-expressed genes were identified in the red module and 274 in the green module. Of these, 66 genes from the red module and 27 from the green module overlapped with the DEGs identified in the GSE6740 dataset (left panel of Fig 4). These overlapping genes were designated as differentially co-expressed genes.

**Fig 4.**
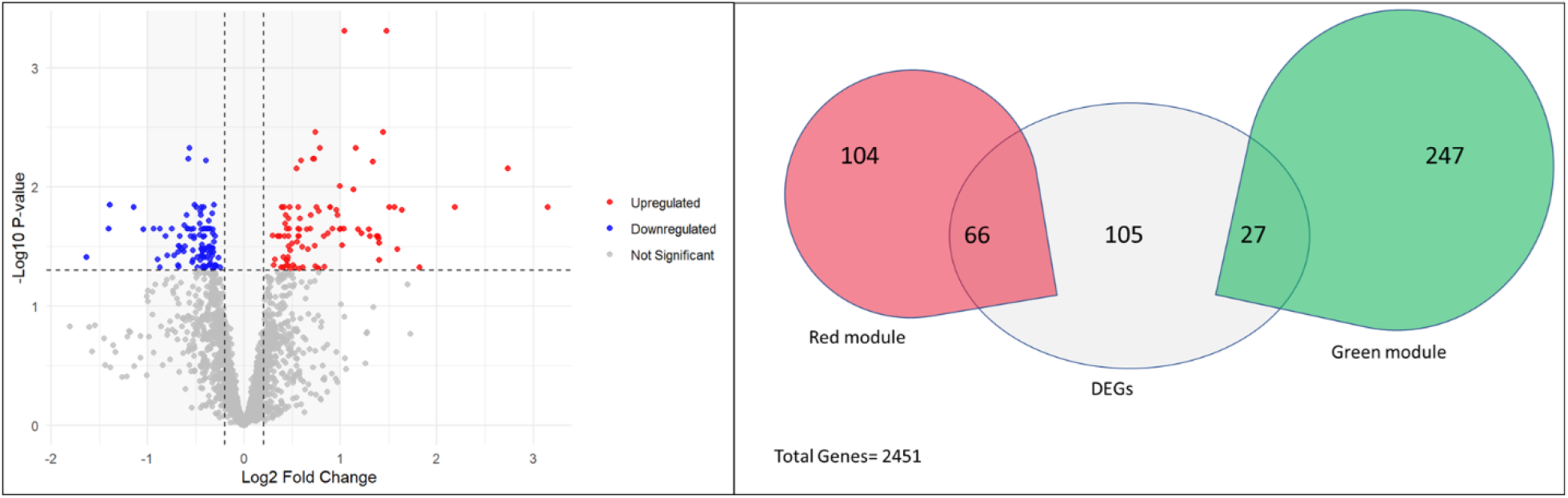
Differentially expressed genes (DEGs) from GSE6740 dataset and intersection with yellow module genes.

**Fig 5.**
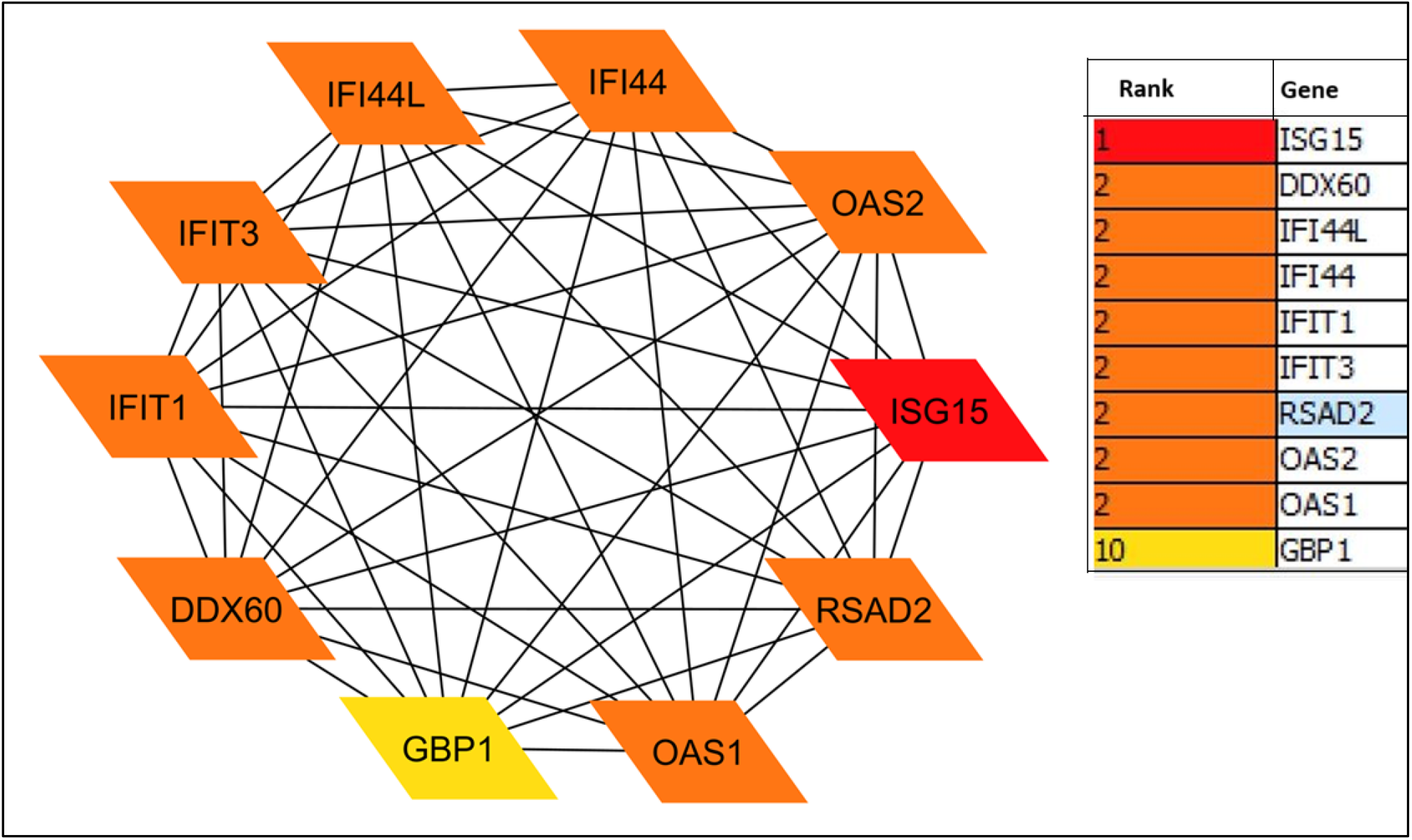
Ten hub genes with ranks from PPI network analysis.

#### Gene Ontology (GO) and KEGG Pathway Analysis

A total of 80 unique differentially co-expressed genes (from both the red and green modules) were used for Gene Ontology (GO) and KEGG pathway enrichment analysis. These 80 genes were selected from 93 by removing duplicate genes that were co-expressed in both modules. The enrichment results revealed highly significant GO terms, indicating strong activation of antiviral defense mechanisms in HIV-infected individuals (S1 Tables 1). Notably, the enrichment of negative regulators of viral processes suggests upregulation of host genes involved in controlling HIV replication. Key upregulated genes include ISG15, IFIT1, OAS1/2/3, RSAD2, MX1, TRIM22, PLSCR1, and BST2, all of which are interferon-inducible genes with established anti-HIV activity (S1 Fig 3).

KEGG pathway enriched reflect the activation of shared antiviral immune responses-such as interferon-stimulated genes (ISGs), inflammatory cytokines, and cellular defense mechanisms (S1 Table 2). Although these pathways are labeled for different viruses (e.g., Influenza A, Hepatitis C), the shared components of host antiviral defense (e.g., IFITs, ISG15, MX1, OAS1-3) are likely upregulated in HIV infection as well. This suggests cross-pathogen immune activation—a signature of chronic HIV infection due to ongoing immune stimulation and inflammation. (S1 Fig 4).

#### PPI Network Analysis and Hub Gene Identification

PPI network analysis was performed by uploading selected 80 genes (both red and green) on the STRING website. Then the PPI network was imported in Cytoscape for hub gene screening (S1 Fig 6). Using MCC algorithm top ten genes with the highest scores were defined as Hub genes, including: DDX60, IFI44L, IFI44, IFIT1, IFIT3, RSAD2, OAS2, OAS1, ISG15, and GBP1 (Fig 4).

### Identification of the Optimal HIV Diagnostic Model

Machine learning models were developed using clinical information (such as age and treatment group) along with the expression profiles of ten hub genes to identify the most effective HIV diagnostic model. Among the six models evaluated-Logistic Regression, GLMNET, SVM, kNN, Decision Tree, and Random Forest - GLMNET emerged as the best-performing model overall. It achieved the highest mean ROC (0.99), indicating excellent discriminative ability, and also recorded the highest mean accuracy (0.92) across 30 resamples (Table 1; Figs 7–8). In terms of specificity (0.92) and kappa (0.72), GLMNET outperformed all other models, reflecting both strong classification performance and substantial agreement beyond chance. While its sensitivity (0.733) was slightly lower than that of Logistic Regression (0.97), GLMNET maintained a better balance between sensitivity and specificity, making it a more reliable model overall. Although Random Forest and Decision Tree also performed well, GLMNET provided the most consistent and robust performance across all key evaluation metrics, making it the most suitable model for the dataset. The best-performing GLMNET model was obtained with the hyperparameters alpha = 0 and lambda = 0.29, identified through 10-fold cross-validation using a grid search over a range of alpha and lambda values. This optimal combination was selected based on the highest cross-validated accuracy from the training results.

**Table 1.** Average accuracy, Kappa, ROC (AUC), sensitivity, and specificity for each model estimated using 10-fold cross-validation.

| Model | Accuracy | Kappa | ROC | Sensitivity | Specificity |
| --- | --- | --- | --- | --- | --- |
| RandomForest | 0.90 | 0.67 | 0.94 | 0.70 | 0.93 |
| DecisionTree | 0.91 | 0.68 | 0.86 | 0.76 | 0.96 |
| kNN | 0.83 | 0.56 | 0.88 | 0.70 | 0.86 |
| GLMNET | 0.92 | 0.72 | 0.99 | 0.73 | 0.92 |
| SVM | 0.89 | 0.62 | 0.77 | 0.43 | 0.92 |
| Logistic | 0.83 | 0.70 | 0.91 | 0.97 | 0.78 |

**Fig 6.**
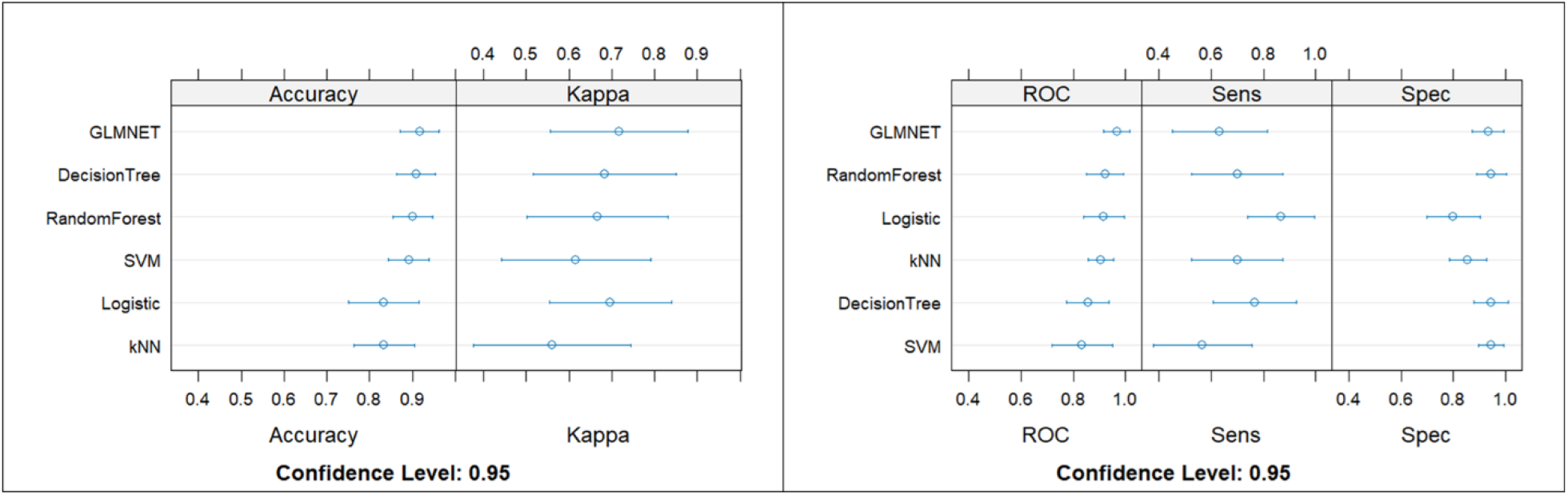
Average accuracy, Kappa, ROC (AUC), sensitivity (Sens), and specificity (Spac) for each model, along with 95% confidence intervals, estimated using 10-fold cross-validation.

**Fig 7.**
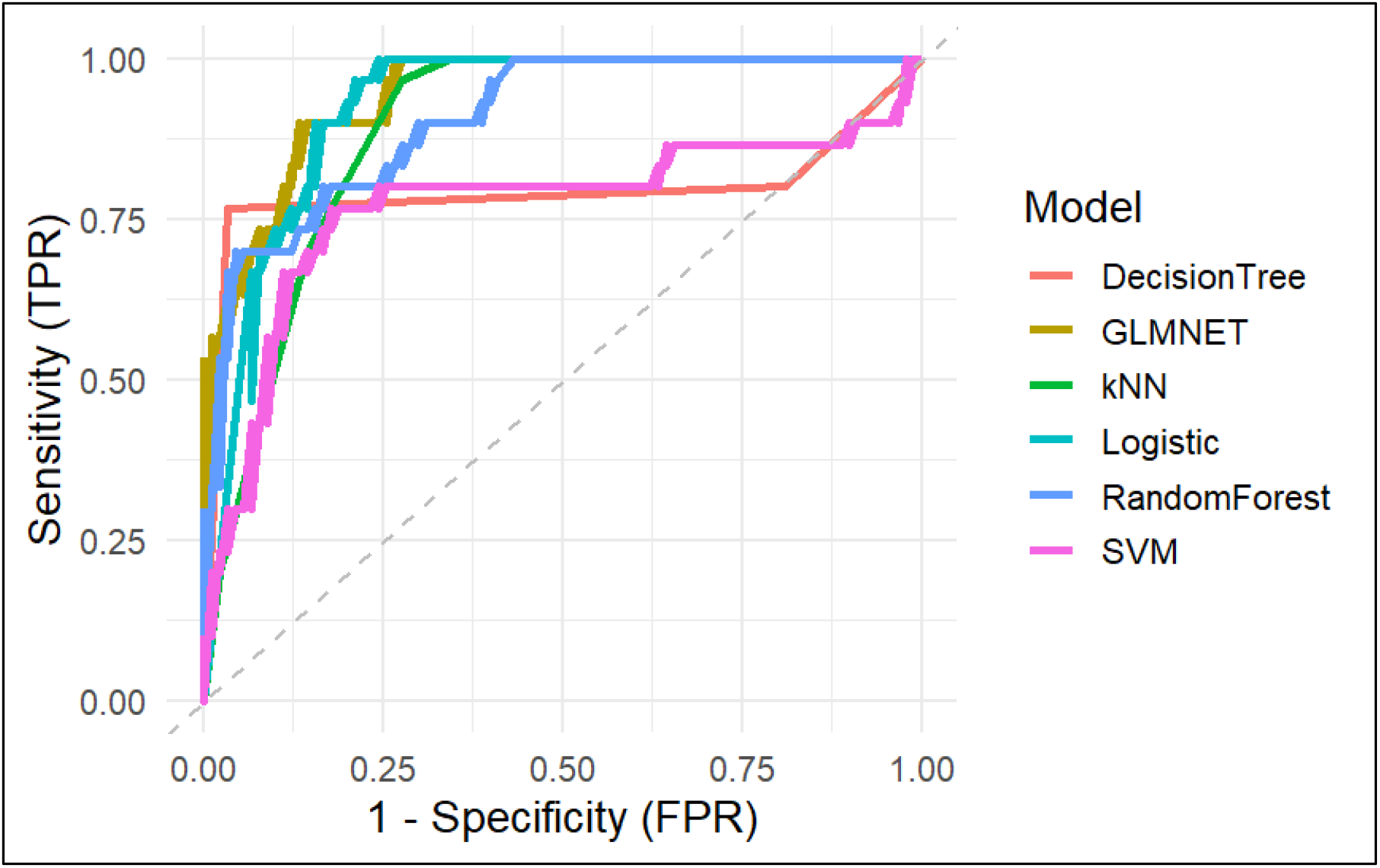
ROC curve comparison among selected machine learning models.

### Interpretation and Visualization of the HIV Diagnostic Model

This SHAP analysis provided insights into the key drivers of the model’s predictions. The model’s average predicted probability was 0.75, while the analyzed sample showed a higher prediction of 0.90 (S1 Fig 6(a-f)), indicating a positive deviation primarily influenced by specific features.

The most impactful features were GBP1, ISG15, OAS2, OAS1, and DDX60, which consistently pushed predictions higher. GBP1 emerged as the strongest contributor overall. Age also influenced predictions both positively and negatively, while the treatment group had lowest impact (Fig 8 (a-b), S1 Fig 6 (a-f)).

**Fig 8.**
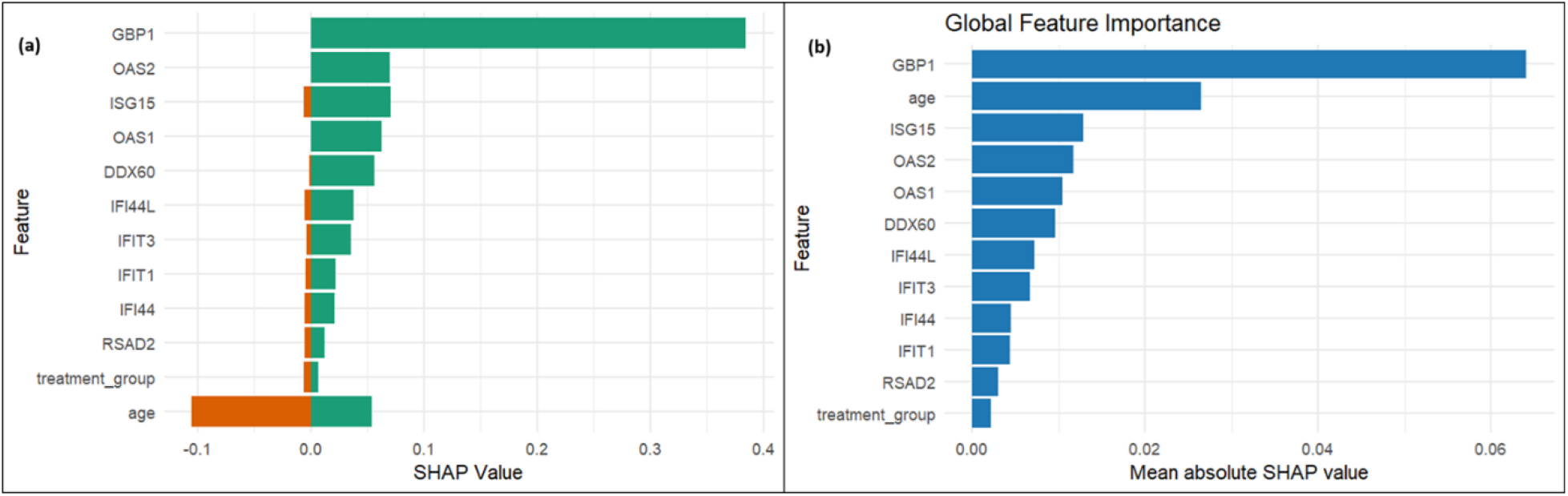
(a) Summary of SHAP Values Across Features. (b) Global feature importance based on absolute shape value.

## Discussion

This study presents a novel integrative framework that combines immune cell–specific transcriptomic data with interpretable machine learning to identify robust and biologically relevant biomarkers for HIV infection. Traditional clinical biomarkers, such as CD4+ T cell count and viral load, while essential, do not fully capture the molecular complexity of HIV pathogenesis [3]. Our findings emphasize that transcriptomic signatures derived from specific immune cells, analyzed through systems biology and machine learning pipelines, can provide deeper insights into host– virus interactions [23, 24].

The WGCNA identified co-expression modules significantly associated with HIV status, particularly the red and green modules, which showed the strongest correlation. Integration with differentially expressed genes (DEGs) and protein–protein interaction (PPI) networks allowed us to isolate hub genes that are not only statistically significant but also biologically central to the underlying network, potentially reflecting key drivers of HIV-related immune dysregulation. These genes enriched Gene Ontology (GO) terms related to immune response, viral processes, and apoptosis, aligning with known mechanisms of HIV pathology [25].

Among the various machine learning (ML) classifiers developed, the GLMNET model-optimized through hyperparameter tuning-achieved the best diagnostic performance, with a high area under the curve (AUC). This is consistent with prior findings that regularized models like GLMNET are well-suited for high-dimensional omics data [26, 27]. Importantly, SHapley Additive exPlanations (SHAP) values enhanced model interpretability by identifying GBP1, ISG15, OAS2, OAS1, and DDX60 as the most impactful genes for differentiating HIV-positive from HIV-negative individuals. Notably, GBP1 has also been reported as a novel biomarker for chronic inflammatory diseases [28], while previous studies suggest that ISG15 and OAS2 may play important roles in the anti-HIV-1 response of CD4+ and CD8+ T cells [29].

Our study underscores the utility of combining biologically informed feature selection methods with interpretable AI to enhance the diagnostic landscape for HIV. By using only peripheral immune cell gene expression profiles, we offer a minimally invasive yet highly informative approach to support early HIV diagnosis. Additionally, the identified hub genes may serve as potential targets for future therapeutic research or as candidates for longitudinal monitoring of treatment response.

However, the study has limitations. First, the findings are based on a single dataset (GSE6740) with a relatively small sample size, which may limit generalizability. Future studies should validate the model on larger and more diverse cohorts. Second, while SHAP provides model interpretability, experimental validation of the selected biomarkers is necessary to establish causal relevance. Lastly, the transcriptomic data represents a static snapshot; integrating longitudinal data or single-cell sequencing could further enrich the diagnostic model and improve precision.

In conclusion, our study demonstrates that integrating transcriptomic profiling with machine learning and explainable AI can lead to the discovery of clinically meaningful and interpretable biomarkers for HIV diagnosis. This approach has the potential to complement existing diagnostic tools and contribute to more personalized and timely HIV care strategies.

## Data availability statement

Data file is publicly available from the Gene Expression Omnibus (GEO) database (https://www.ncbi.nlm.nih.gov/gds). All code files are available from Open Science Framework repository at https://doi.org/10.17605/OSF.IO/WMQ7K.

## S1: supportive information

## Notes

### Competing Interest Statement

The authors have declared no competing interest.

